# Mobilization of genes encoding potential PFAS-degradation enzymes and positive selection in cyanobacteria

**DOI:** 10.1101/2025.07.23.666183

**Authors:** Asal Forouzandeh, Malte Storm Lau Schlosser, Annaliese Nan Vernon, Tue Kjærgaard Nielsen

## Abstract

Per- and polyfluoroalkyl substances (PFAS) are persistent environmental pollutants due to their strong carbon-fluorine bonds and the toxicity of released fluoride from degradation, posing significant challenges for bioremediation. While microbial defluorination of PFAS has been described, the enzymatic mechanisms and evolutionary selection driving this capability is poorly understood. In this study, we screened all complete bacterial genomes in the NCBI RefSeq database for homologs of 43 candidate defluorinating enzymes, focusing on their genetic context and association with mobile genetic elements. Our analysis revealed that fluoroacetate dehalogenases and haloacid dehalogenases are among the most frequently mobilized enzyme classes, suggesting positive selection. Notably, a conserved gene encoding a fluoroacetate dehalogenase homolog in *Microcystis* cyanobacteria is consistently flanked by genes related to stress response, photosynthesis, and toxin-antitoxin systems, suggesting a potential adaption to alleviate PFAS toxicity. We propose that positive selection for PFAS degradation may be driven by mitigation of physiological stress rather than metabolic gain. These findings highlight the importance of considering ecological and genetic contexts in the search for effective PFAS-degrading microorganisms.

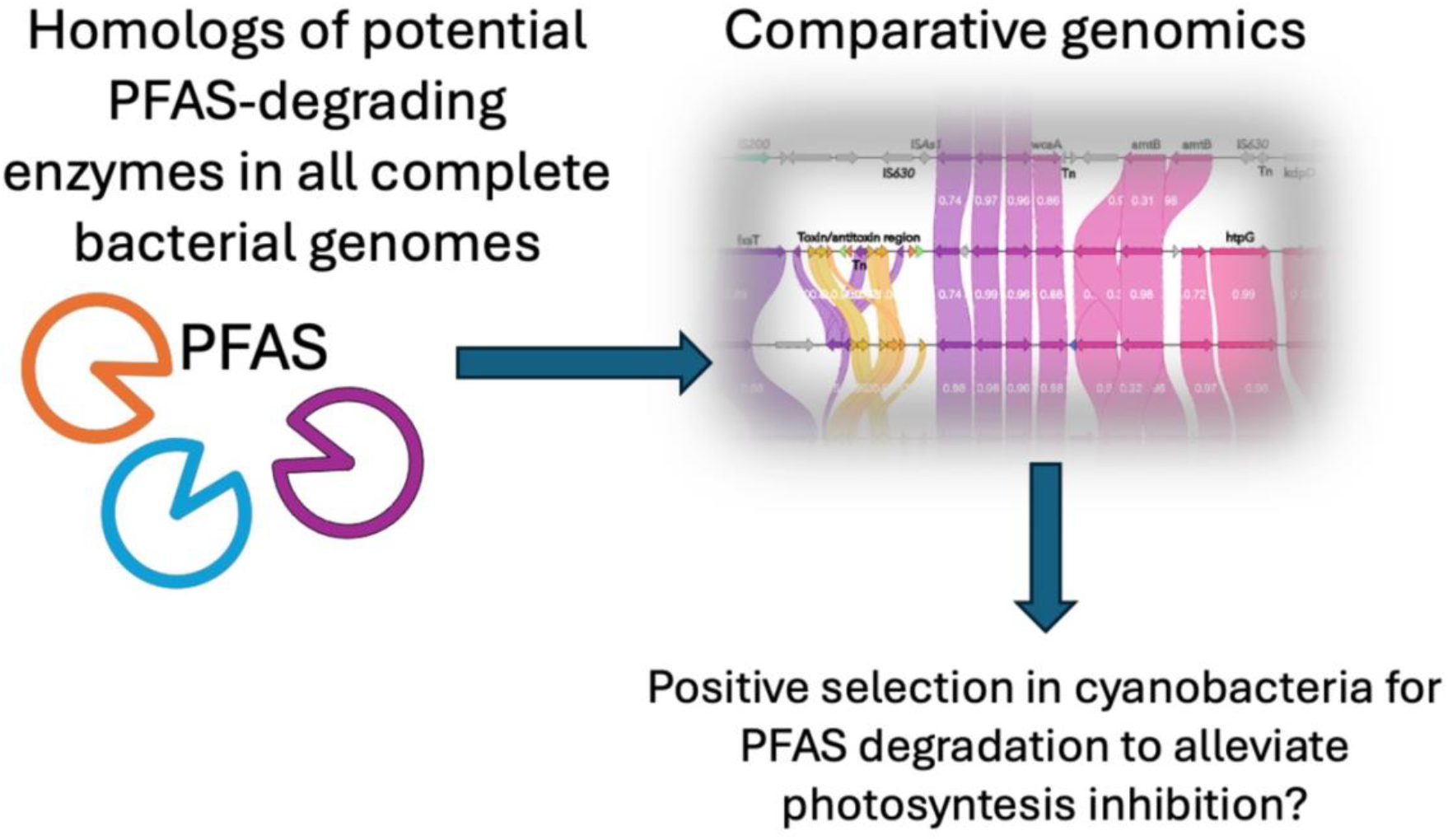

## Introduction

Perfluoroalkyl and polyfluoroalkyl substances (PFAS) are a class of synthetic fluorinated compounds characterized by an aliphatic carbon chain in which some or all of the hydrogen atoms have been substituted with fluorine atoms, typically with the chemical structure C_n_F_2n+1_−R (Wang *et al.* 2017; Presentato *et al.* 2020). The extensive use of PFAS, combined with their inherent resistance to degradation, results in their persistence in both the environment and the human body, where their prolonged biological half-life contributes to bioaccumulation and potential toxicological risks. Epidemiological studies have identified links between exposure to certain PFAS and various health effects, such as altered immune and thyroid function, liver disease, lipid and insulin dysregulation, kidney disease, reproductive and developmental issues, and cancer. Many of these findings are supported by experimental animal data (Fenton *et al.* 2021).

A variety of traditional pollution control methods have been used to tackle PFAS contamination. Bioremediation strategies for fluorinated compounds are still at the beginning of their development and face several hurdles, such as the stability of the carbon-fluorine (C-F) bond and the scarcity of enzymes in nature capable of catalyzing its cleavage (Parsons *et al.* 2008; Liu and Mejia Avendaño 2013). While our understanding of microbial metabolism of fluorinated compounds remains limited, some progress has been made towards microbial biodegradation of highly fluorinated alkyl compounds.

Numerous studies have highlighted the potential of bacteria to defluorinate PFAS (Yi *et al.* 2016; Huang and Jaffé 2019; Shaw *et al.* 2019; Presentato *et al.* 2020; Marchetto *et al.* 2021). A wide variety of bacteria have been investigated under various experimental conditions. For instance, *Acidimicrobium* sp. strain A6 has demonstrated the capability to degrade >60 % of the recalcitrant PFOA and PFOS over a 100-day period while reducing iron and utilizing ammonium or hydrogen as electron donors (Huang and Jaffé 2019). Another study found that *Labrys portucalenis* F11, an aerobic bacterium isolated from a contaminated site, could degrade 90% of PFOS, 58% of 5:3 FTCA, and 21% of 6:2 FTS after 100 days of incubation, with an increase in free fluoride concentration and detectection of shorter chain PFAS as degradation products (Wijayahena *et al.* 2025). Similarly, *Pseudomonas parafulva* has been reported to degrade PFOA by utilizing it as a carbon source (Yi *et al.* 2016). Additionally, *Gordonia sp.* strain NB4-1Y has shown the ability to degrade 6:2 fluorotelomer sulfonamidoalkyl betaine, along with its potential degradation intermediate (Shaw *et al.* 2019). Several other studies have reported perfluoroalkyl acid (PFAA) degradation in microbial consortia and enrichment cultures, although the specific enzymes responsible for defluorination remain unidentified in most cases (Smorada, Sima and Jaffé 2024).

### Proposed enzymes for defluorination of PFAS

Fluoroacetate, a naturally occurring monofluorinated compound produced by certain plants and bacteria (Murphy, Schaffrath and O’Hagan 2003), has been shown to be degraded by bacterial fluoroacetate dehalogenase enzymes (Leong *et al.* 2017). Multiple classes of enzymes have been proposed to be involved in degradation of PFAS, including cytochrome P450s, laccases, peroxygenases, hydratases, monooxygenases, dioxygenases, dehydrogenases, reductases, and the fluoroacetate-, haloacid-, and haloalkane dehalogenases. Their catabolic activity on PFAS range from speculative (Chetverikov *et al.* 2023) to detectable yet insignificant release of fluoride (Harris *et al.* 2022). As such, very few enzymes with efficient PFAS degradation capability have been described yet. Of the proposed classes of enzymes, undiscovered variants of some of the dehalogenases may hold promise for PFAS degradation, specifically the perfluorinated alkyl acids (PFAA), including PFOA and PFOS (Hu and Scott 2024; Wackett 2025). The haloacid dehalogenase DeHa1, cloned from *Delftia acidovorans* into *E. coli*, showed detectable but not significant defluorination of PFOA (Harris *et al.* 2022). A reductive dehalogenase RdhA from anaerobic *Acidimicrobium* sp. A6 demonstrated defluorination of PFAAs coupled to the Feammox reaction (Jaffé *et al.* 2024). *Pseudomonas mosselii* can utilize PFOA and other perfluoroalkyl carboxylic acids (PFCAs) as sole carbon sources, with reductive defluorination at the α-carbon. The researchers speculated that a haloalkane dehalogenase (DhaA) and a haloacetate dehalogenase (DehH1) might be involved (Chetverikov *et al.* 2023). Homologs of fluoroacetate dehalogenases, part of the α/β hydrolase superfamily, are proposed as candidates for PFAS degradation due to their activity on mono-, di-, and trifluorinated α-carbons (Hu and Scott 2024). These enzymes, found in phylogenetically diverse bacteria, include characterized members such as RPA1163, DAR3835, and NOS0089 (Khusnutdinova *et al.* 2023). Other bacterial enzymes potentially involved with defluorination of PFAS include cytochromes P450, peroxidases, dehydrogenases, hydratases, reductases, and dioxygenases acting on aromatic fluorinated compounds (Hu and Scott 2024).

### Potential selection for PFAS degradation

A strong selection for a phenotypic trait can cause the involved genes to be associated with mobile genetic elements (MGEs) that have rearranged genetic elements to confer novel phenotypes. The most obvious case for this is with resistance to antibiotics, where resistance genes mobilized by e.g. plasmids, transposons, and/or integrons are under strong positive selection in human pathogens (Nielsen, Browne and Hansen 2022). Similarly, many xenobiotics can be valuable carbon- and energy sources to bacteria that have adapted to utilize them, often via pathways formed by mobilization of genes encoding catabolic enzymes (Nielsen *et al.* 2013, 2017; Pearce, Oakeshott and Pandey 2015). However, the case of PFAS degradation might pose a different challenge. Most PFAS would serve poorly as carbon- and energy sources, due to the strength of the C-F bonds (Wackett 2021). Furthermore, fluoride anions would be released from enzymatic defluorination, which is likely more toxic to cells than the parent PFAS compound (Stockbridge and Wackett 2024). As such, there is likely no positive selection for the evolution of PFAS-degrading bacteria in most ecological niches. However, recent studies show that PFAS can have specific detrimental effects on bacterial physiology, including disruption of membranes and cytosolic content, induction of oxidative stress, and inhibition of photosynthesis in cyanobacteria (Liu *et al.* 2016; Yang *et al.* 2017; Fitzgerald *et al.* 2018; Fitzgerald, Simcik and Novak 2018; Marchetto *et al.* 2021; Naumann *et al.* 2022; Lindell *et al.* 2024; Bhattacharya *et al.* 2025). As such, it can be hypothesized that any positive selection for PFAS degradation is likely more related to the toxic effects of PFAS, rather than carbon- and energy sources.

In this study, we investigate the genetic context of gene homologs encoding previously hypothesized PFAS-degradation enzymes in all complete bacterial genomes (RefSeq). We specifically investigate MGEs in close proximity to target genes, indicating a potential positive selection for degradation. Concurrent with recent studies suggesting likely enzyme candidates (Hu and Scott 2024; Wackett 2025), we find that genes encoding fluoroacetate dehalogenases (FAcD) and haloacid dehalogenases (HAD) are the most mobilized, although they are rare in the RefSeq bacterial genome database. We further investigate the conserved genetic organization around a FAcD in cyanobacterial *Microcystis* chromosomes, which includes several genes related to cellular stress, genetic stability (toxin-antitoxin modules), and photosynthesis, suggesting a coupling between these functions and potential PFAS degradation. In light of these results, we propose that future efforts to find PFAS-degrading microorganisms should consider the reason and possible evolutionary selection for their removal.

## Materials and Methods

### Screening for homologs of potential defluorinating enzymes

All scripts and data files, except third party databases (e.g. RefSeq genomes), are available from Github: https://github.com/tueknielsen/Defluorination_genes/.

Complete bacterial genomes were downloaded from the NCBI RefSeq database in June 2024 using the ‘ncbi-genome-download’ tool v.0.3.3 (Blin 2023). Taxonomic and replicon information for each hit was extracted by matching accession numbers to the prokaryotes overview files (https://ftp.ncbi.nlm.nih.gov/genomes/GENOME_REPORTS/prokaryotes.txt and https://ftp.ncbi.nlm.nih.gov/genomes/GENOME_REPORTS/plasmids.txt), with adjustments for discrepancies in accession formats. All genome FASTA files were concatenated into a single file for downstream analysis. Gene homologs encoding potential defluorinating enzymes were identified using BLASTX in DIAMOND v.2.1.8.162 (Buchfink, Xie and Huson 2015) against a custom database of 43 potential defluorinating enzyme sequences (Supplementary file target_enzymes.faa and Supplementary data). This custom database includes enzymes shown to defluorinate fluorinated compounds (including non-PFAS e.g. fluorobenzoate) and enzymes only suspected to be involved in degradation of fluorinated compounds based on e.g. transcriptomic data. Hits were filtered based on identity (≥60%), subject coverage (≥80%), and E-value (≤10E-10). Sequence coordinates of target gene hits were extracted and adjusted to include 12,170 bp of flanking regions on both sides of gene hits. This distance matches the previously determined average size of unit and composite transposons (Nielsen, Browne and Hansen 2022).

Nucleotide sequences of hit regions were extracted as fasta files for further processing. The proximity of target genes to insertion sequence (IS) elements was determined by running DIAMOND BLASTX against the ISfinder database (Siguier *et al.* 2006), as implemented in TnCentral (Ross *et al.* 2021) updated May 2024. The closest IS elements upstream and downstream of each target gene, encoding potential defluorinating enzymes, were identified and distances were calculated.

### Clustering genetic regions and statistical analyses

Target gene regions were clustered using USEARCH v11.0.667 (Edgar 2010) at 99% identity and 90% query/subject coverage to identify Clustered Hit Loci (CHLs). A compression ratio was calculated, expressed as CHLs divided by number of unclustered hit regions. Integron detection was performed using IntegronFinder v.2.0 (Néron *et al.* 2022) on clustered loci. Integron-associated genes were identified and annotated, but no target defluorination genes were found within integrons. Therefore, the presence of genes encoding potential defluorinating enzymes within integrons was not considered in downstream analyses.

RStudio was used for statistical analyses and data visualization. The following R packages were used: pheatmap (Kolde 2025), dplyr (Wickham 2015), reshape2 (Wickham 2007), vegan (Oksanen 2015), tidyr (Wickham 2024), and ggplot2 (Wickham 2016). Taxonomic distribution across distinct genera (quantified by Simpson’s diversity index), IS proximity ratios, plasmid/chromosome rates, and clustering compression rates were calculated for each CHL. Results from all analyses were integrated into a comprehensive dataset. A unifying MOBscale (Nielsen, Browne and Hansen 2022) was calculated as the mean of the Simpson index, IS proximity ratio, and replicon type ratio, which are all on a 0 to 1 scale, with 1 representing the highest mobilization and 0 no mobilization of a given CHL with a target gene. Comparative genomics plots were made with Clinker v.0.0.31 (Gilchrist and Chooi 2021) using genbank files for CHLs made with Bakta v.1.11.0 (Schwengers *et al.* 2021). Gene features in Clinker plots were manually inspected and curated using online BLASTP (Altschul *et al.* 1990).

## Results and Discussion

### Sequence comparison of defluorination enzymes

Amino acid sequences of 43 target enzymes (Supplementary file target_enzymes.faa) were pairwise compared for amino acid similarity with BLASTP and subsequent hierarchical clustering (Supplementary Fig. S1). The included enzymes represent multiple classes and superfamilies (Supplementary Table 1), showing distinct groupings based on sequence similarity. Three groupings occur with three oxidoreductases, six fluoroacetate dehalogenases (FAcDs), and two groups of three more dissimilar haloacid dehalogenases (HADs) (Supplementary Fig. S1). The included enzymes have either proven defluorination activity or indications of activity based on transcriptomics or other inconclusive results (Supplementary Table 1). The included dioxygenases, hydrolases, reductases, dehydrogenases, cytochrome P450, peroxygenase, and haloalkane dehalogenase have either activity on non-PFAS aromatic fluorinated compounds (e.g. phenols and benzoates) or, in the case of monooxygenases and oxidoreductases, only speculative activity on PFAS based on transcriptomics (Supplementary Table 1). We therefore focus mostly on HADs and FAcDs, as homologs of these are more likely to activity on perfluorinated alkyl acids (Hu and Scott 2024; Wackett 2025).

The six included FAcDs share the previously described catalytic triad Asp110-His280-Asp134 (RPA1163 numbering; not shown) (Khusnutdinova *et al.* 2023; Hu and Scott 2024). Both HADs and FAcDs are homodimer proteins and some have defluorination activity on mono- and difluoroacetate but not yet shown for other fluorinated compounds (Khusnutdinova *et al.* 2023). However, the HAD RdhA (QAX87819.1) is likely involved in PFOA and PFOS degradation in the anaerobic *Acidimicrobium* sp. A6, based on loss of defluorination on these substrates in a RdhA knockout mutant (Jaffé *et al.* 2024). Unfortunately, enzymes QAX87819.1 and WP_011137954.1 representing RdhA from *Acidimicrobium* sp. A6 and DeHaII haloacid dehalogenase type II from *Delftia acidovorans*, respectively, were not found in any complete genomes and were not included in downstream analyses.

### Search results and clustering

A search for homologs of defluorinating enzymes using DIAMOND BLASTX against all complete bacterial genomes from the RefSeq database yielded 15,908 hits with a minimum E-value of 10E-10, similarity of 60% (ID), and query coverage of 80% (Fig. 1). The median percent ID were highest for those enzymes with very few hits, but also high for the very abundant haloacid dehalogenase AAG04199.1. In this study, we do not distinguish between the origin of enzyme homology (paralogous or orthologous), but all enzymes passing the above filters are assumed to be isofunctional, although this might not always be true. The number of hits ranged from 3,922 (WP_053776268.1) to 1 (WP_053777670.1, DAR3835, YP_00337005.1). These extremes represent several factors, including the skewness towards easily culturable bacteria in the RefSeq complete genomes, the rarity of the query enzymes in bacteria, and database biases towardshuman associated bacteria. The HAD AAG04199.1 is for example found 877 times in the database but only in 21 distinct genera, of which *Pseudomonas aeruginosa* makes up 94.4% with near identical sequences. The FAcD DeHa4 is the most abundant of this enzyme class with 1,060 unclustered hits and occurs in 57 distinct genera, mainly in *Burkholderia* (41.0%) and *Pseudomonas* (21.1%). The query DeHa4 enzyme was originally described in a *Delftia* (Harris *et al.* 2022), but this genus only constitutes 2.3% of the genomes with homologs. For the dehydrogenase WP_053776268.1, the 3,923 hits are distributed across 148 genera, with *Bacillus, Streptomyces*, and *Mycobacterium* making up 25%, 14,8%, and 14.6%, respectively. Besides reflecting biological properties of the enzymes, such as their importance to essential/core functions, this shows the biases towards some groups of bacteria that skew the results of some enzymes more than others. To compensate, clustering of identical flanking regions to Clustered Hit Loci (CHLs) was performed.

**Fig. 1.**
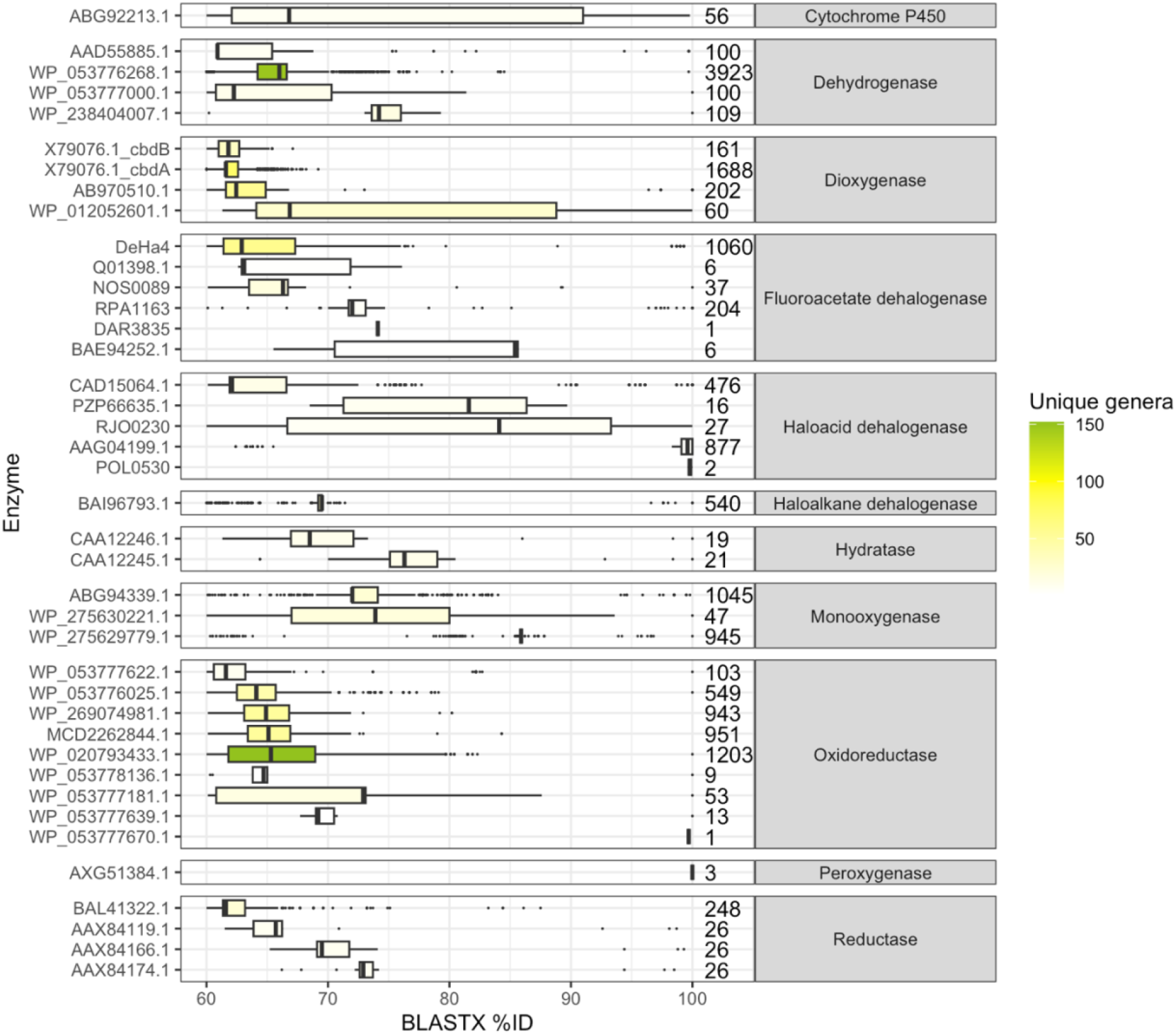
The % identity of DIAMOND BLASTX hits for each defluorinating enzyme. Numbers to the right indicate the number of unclustered hits.

From all BLASTX hits, 12,170 bases were extracted flanking the hit, corresponding to the average size of composite and unit transposons, as previously described (Nielsen, Browne and Hansen 2022). The total hits (15,908) were clustered to Clustered Hit Loci (CHLs) that are at least 99% similar (nucleotide) over at least 90% of the sequence length. The compression rate for each enzyme was calculated (Supplementary Fig. S2), where 100% indicates that all hit regions were compressed into a single CHL. This is the case when all gene matches plus flanking regions are identical or almost identical in all genomes where it is found. Conversely, a lower compression rate shows that a given gene is found in diverse contexts across different genomes. The highest compression rates are associated with genes that have few matches in RefSeq complete genomes (BAE9452.1, DAR3835, POL0530, Q01398.1, WP_053777670.1). These are completely compressed to a single CHL, showing that they are found in highly similar (or identical) contexts across different genomes. Only DAR3835 and WP_053777670.1 were found in only one genome. The most abundant gene was WP_053776268.1, encoding a NAD(P)-dependent alcohol dehydrogenase, which was found in 3,923 distinct genomes that could be clustered to 2,140 unique CHLs (Fig. 1, Supplementary Fig. S2). The lowest compression rates, indicating the most diverse genetic contexts, were found for BAI96793.1, WP_275629779.1, AAG04199.1, encoding 1,3,4,6-tetrachloro-1,4-cyclohexadiene hydrolase LinB, alkane 1-monooxygenase, and probable haloacid dehalogenase, respectively. The highly abundant HAD enzyme AAG04199.1 (n=877; Fig. 1) had a median similarity of 99.6% and was found in *Pseudomonas aeruginosa* in 94.4% of cases. However, its low compression rate of 19.2% shows that the flanking context of the hits is very varied, indicating a high selection for sequence conservation of the enzyme itself but low conservation of the genetic synteny. For the likewise very abundant dehydrogenase WP_053776268.1, the 3,923 hits were compressed to 2,140 CHLs for a compression rate of 54.6%, indicating intermediate conservation of genetic synteny, which is also reflected by a low median similarity of 66% (Fig. 1). The enzymes with fewer hits are generally also more compressed by clustering, indicating low abundance of these enzymes in the database with a conserved genetic synteny (Supplementary Fig. S2).

### Some homologs of defluorinating dehalogenases are highly mobilized

The MOBscale ratio for each enzyme is calculated as the average of the three mobilization parameters: association with IS elements, plasmid location, and Simpson diversity index. As described above, we focus mostly on HADs and FAcDs as these are proposed as candidates for degradation of perfluorinated alkyl acids (Hu and Scott 2024; Wackett 2025). Aside from these, the gene encoding the alpha-subunit of toluene dioxygenase (TodA; WP_012052601.1) is highly mobilized with a MOBscale score 0f 0.51 (Fig. 2). This enzyme can degrade the fluorinated aromatic compound 2,2-fluoro-1,3-benzodioxole (DFBD) (Bygd *et al.* 2021) but not alkyl substrates.

**Fig. 2.**
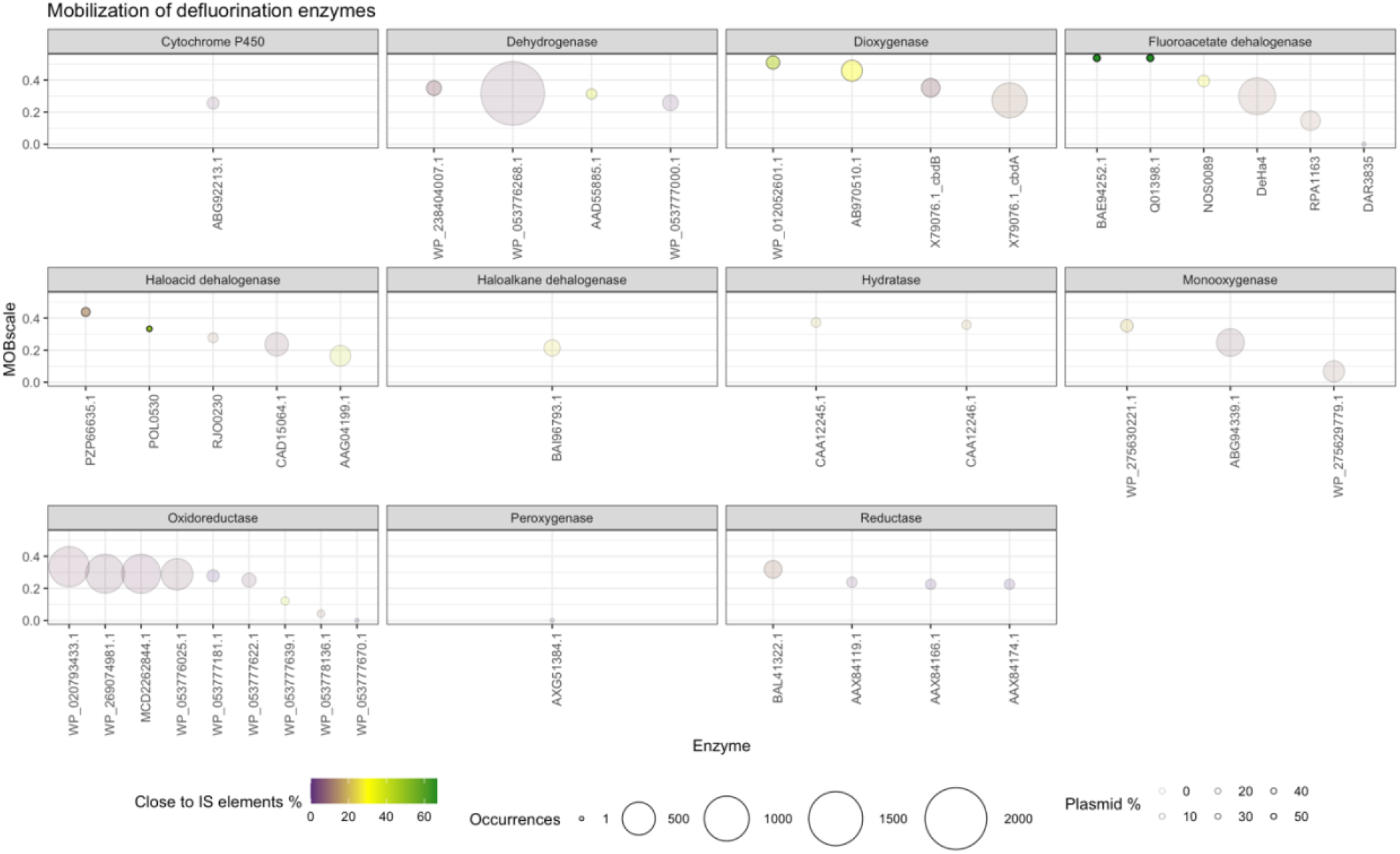
MOBscale and individual mobilization parameters for CHLs of each included gene. The color indicates how often a specific gene is found near IS elements. The size indicates how many CHLs are present for a given gene with more CHLs representing larger genetic diversity. The transparency (alpha) indicates how often CHLs of a given gene are found on plasmids, with more solid circles representing a higher frequency on plasmids.

The most mobilized FAcD-encoding genes are BAE94252.1 and Q01398.1 (Fig. 2), representing DehH fac-dex fluoroacetate dehalogenase from *Burkholderia* sp. FA1 and fluoroacetate dehalogenase H-1 from *Delftia acidovorans,* respectively. The latter has been described as a HAD but contains the FAcD-characteristic conserved catalytic triad domain (not shown) (Khusnutdinova *et al.* 2023) and is more similar in sequence to other FAcDs than HADs (Supplementary Fig. S1). The DehH and H-1 enzymes are 63.3% similar (Supplementary Fig. S1) and match the same six database sequence entries, although with different similarities. The RefSeq complete genomes matches are from two unique genera (*Mesorhizobium* and *Ralstonia*), occur in proximity to IS elements in 4/6 CHLs, and are located on plasmids in 3/6 CHLs.

Homologs of the more abundant, but less mobilized, FAcD NOS0089 are found in 37 RefSeq entries (35 CHLs) with an IS association in 31.42% and plasmid association in 2.7% of regions (Fig. 2). It is found in 18 unique genera, with highest prevalence in *Microcystic, Nostoc*, and *Skermanella*. As described above, the most frequent FAcD homologs were those of DeHa4, with 647 CHLs. They are rarely found on plasmids (0.28%) but often near IS elements (13.1%). The occurrence in 57 distinct genera results in a high Simpson index of 0.78. The FAcD RPA1163 is also abundant with 145 CHLs, distributed across only 5 unique genera, for a very low Simpson diversity index. It is mostly found in *Bradyrhizobium* (82.4%) and *Rhodopseudomonas* (8.33%). It has no occurrences on plasmids but is associated with IS elements in 13.10% of chromosomes. This indicates that RPA1163 is either part of an IS element hotspot chromosomal region in *Bradyrhizobium* or that it is mobilized through other means than plasmids among members of *Bradyrhizobium* (e.g. ICEs that are not included in the MOBscale). The same is likely the case for the FAcD DeHa4 but mainly in *Burkholderia* and *Pseudomonas*, as described above.

The HADs PZP66635.1 from *Delftia acidovorans* and POL0530 from *Polaromonas* sp. are also mobilized in some genomes with MOBscale ratios of 0.44 and 0.33, respectively (Fig. 2). PZP66635.1 is the most abundant of the two (16 hits clustered to 14 CHLs) and is widely distributed to 8 distinct genera (*Achromobacter, Bosea, Bradyrhizobium, Cupriavidus, Ensifer, Martelella, Phreatobacter,* and *Sinorhizobium*). It is associated with IS elements in 14.29% of CHLs and occur on plasmids in 31.25% of entries.

A single homologous sequence to FAcD DAR3835 (74.1% ID) was matched in the RefSeq complete genomes database and is found on the chromosome of *Sulfuritalea hydrogenivorans* sk43H (acc. NZ_AP012547.1). This chemolithoautotrophic facultative anaerobic bacterium was originally isolated from a freshwater lake (Kojima and Fukui 2011) and is involved in anaerobic degradation of aromatic compounds (Sperfeld, Diekert and Studenik 2019). Similarly, DAR3835 was found in the anaerobic *Dechloromonas aromatica* RCB that also degrades aromatic compounds (Salinero *et al.* 2009). The structure of DAR3835 was resolved and high defluorination activity was demonstrated, when expressed in *E. coli*, on difluoroacetate and limited activity on 2,2-difluoropropionic acid and 5,5,5-trifluoropentanoic acid (Khusnutdinova *et al.* 2023). These promising results show that homologs of DAR3835 and similar may have activity on PFAS, although our results show that homologs of the enzyme are rare.

### A NOS0089 homolog is flanked by IS elements, and photosynthesis- and stress-related genes

A conserved homolog of FAcD NOS0089 is found on the chromosomes of 14 different *Microcystis* cyanobacteria (Fig. 3). The genetic context of these FAcD includes several genes associated with stress tolerance and cell homeostasis, toxin-antitoxin modules, and multiple photosynthesis genes that potentially can compensate for PFAS-mediated photosynthesis inhibition. Some PFAS have been shown to inhibit photosynthesis, increase cell permeability, and induce oxidative stress in cyanobacteria in compound- and species-specific ways (Marchetto *et al.* 2021; Bhattacharya *et al.* 2025; Liao *et al.* 2025), leading to a hypothetical positive selection for degradation. The genetic regions flanking the conserved FAcD is also characterized by multiple IS elements and transposases, indicating genetic rearrangements, positive selection, and potential for horizontal gene transfer. In all chromosomes, two genes encoding acetoin utilization protein (*acuC*) and a coenzyme F420-reducing hydrogenase (F420) are sitting upstream of the NOS0089 FAcD homolog. Their link to potential PFAS degradation is not clear, however. A conserved *wcaA* gene encoding a glycosyltransferase, likely involved in cell wall synthesis or biofilm formation, is downstream of the FAcD gene in some *Microcystis* (Fig. 3). Some PFAS have been shown to accumulate in- and damage bacterial cell membranes and increase cell permeability (Liu *et al.* 2016; Yang *et al.* 2017; Fitzgerald *et al.* 2018; Fitzgerald, Simcik and Novak 2018). In *E. coli*, PFAS exposure induces differential expression of many genes, including some involved in membrane stress response (Wintenberg, Vasilyeva and Schaffter 2025), while other bacteria display inhibition of genes related to cell wall biogenesis (Qiao *et al.* 2018). Finally, PFAS exposure can lead to increased extracellular polysaccharide and biofilm formation (Weathers, Higgins and Sharp 2015). Multiple other genes hypothetically linked to cell homeostasis, in response to PFAS exposure and increased cell permeability, are conserved in the vicinity of the FAcD genes, including *amtB, tauE,* and *kdpD* encoding an ammonia channel protein, sulfite exporter, and an osmosensitive kinase sensor regulating the *kdpFABC* potassium channel operon, respectively.

**Fig. 3.**
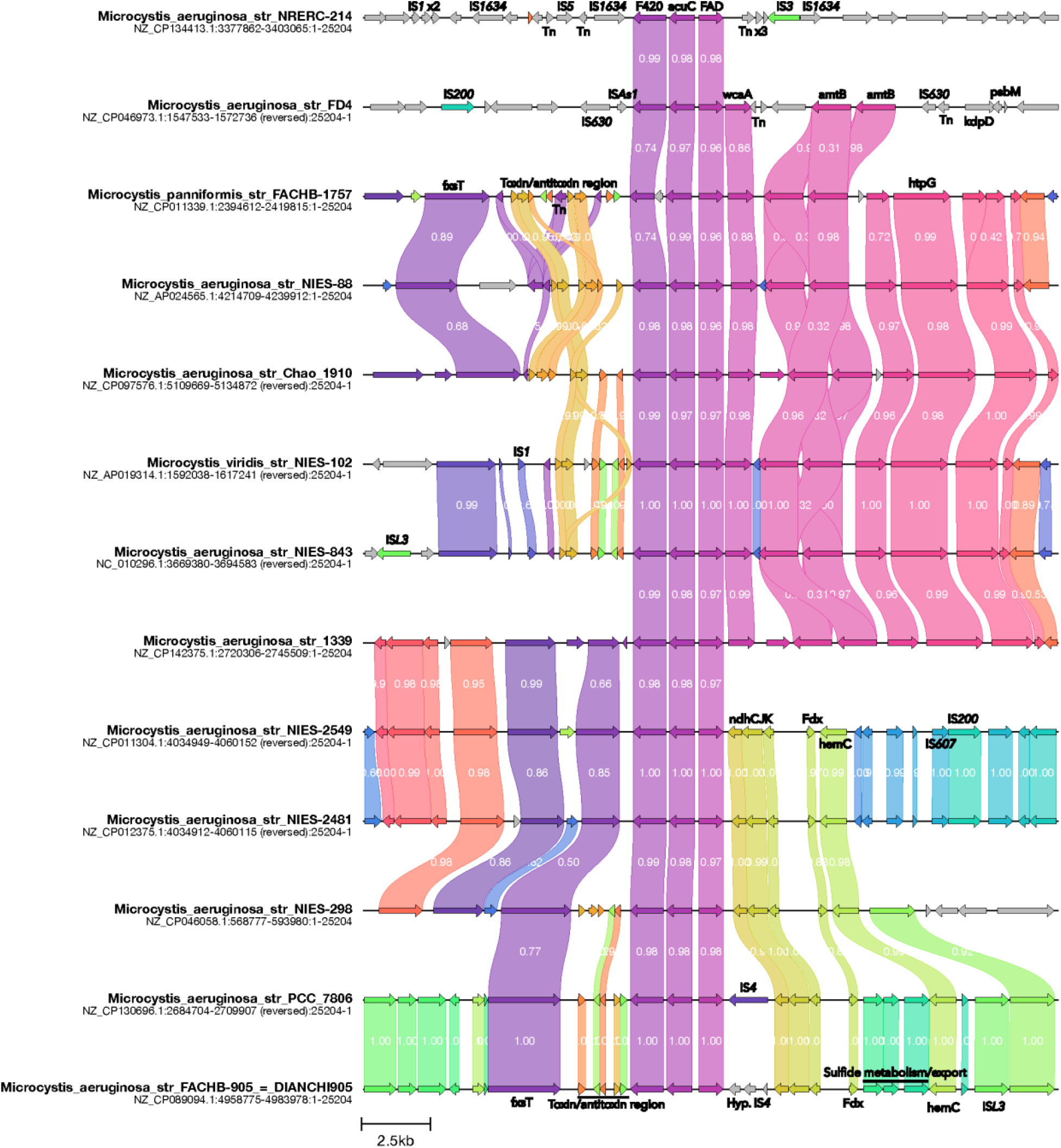
Genetic context of FAcD NOS0089 homologs in M. aeruginosa chromosomes. Numbers in links indicate similarity between amino acid sequences of similar genes. Annotations were manually curated to highlight genes of interest, including IS elements, toxin-antitoxin modules, stress tolerance genes, and photosynthesis-related genes. All instances of M. aeruginosa from complete RefSeq genomes are shown. The figure was produced with Clinker and manually curated with InkScape.

Several variations of a toxin-antitoxin (TA) region occur on 9/14 of the *Microcystis* chromosomes (Fig. 3). TA modules are often associated with selfish MGEs, such as plasmids and transposons, but their function in persistence and stress tolerance have also been speculated, although the latter is disputed (Rosendahl *et al.* 2020). They also act to stabilize nearby genetic elements, including chromosomally integrated MGEs, integrons, and genomic islands (Díaz-Orejas, Espinosa and Yeo 2017). The linkage to FAcD and other genes, indicates a genetic maintenance function and likely positive selection of the genetic organizations. Multiple IS elements occur in the shown regions, which may form transposons that include the TA modules.

Multiple photosynthetic related genes occur in proximity of the FAcD genes (Fig. 3). This is noteworthy since some PFAS have shown to inhibit photosynthesis in cyanobacteria (Bhattacharya *et al.* 2025; Liao *et al.* 2025). PFOA-induced oxidative stress is likely alleviated by overcompensation of the photosystem II (PSII) in both *Microcystis aeruoginosa* (Hu *et al.* 2023) and eukaryotic microalgae (Liu *et al.* 2021, 2025). The *psbM* gene encodes a PSII reaction center protein and Fdx is an electron carrier in photosynthesis (Fig. 3). The *hemC* gene encodes a porphobilinogen deaminase that is part of heme biosynthesis and is possibly linked to chlorophyll synthesis in cyanobacteria. NdhCKJ are structural subunits of the NDH-1 complex that is involved in cyclic-electron-transfer around photosystem I (PSI CET) which is important for alleviating excessive light energy during photosynthesis (Hualing 2022; Bhattacharya *et al.* 2025). In tobacco, knocking out *ndhCKJ* leads to accumulation of reactive oxygen species (Wang *et al.* 2006). The *fsxT* gene encodes a tetratricopeptide repeat (TPR) protein that are involved with proper assembly and function of photosynthetic complexes in cyanobacteria (Kong, Xu and Hu 2003; Klinkert *et al.* 2004).

Taken together, the genetic context of a gene encoding a suspected PFAS-degradation FAcD in multiple *Microcystis* cyanobacteria suggests a selection towards genetic accumulation of functions to counter the toxic effects of PFAS, including cell permeability and photosynthesis inhibition (Fitzgerald, Simcik and Novak 2018; Bhattacharya *et al.* 2025). This genetic organization may be further conserved by the presence of a TA region adjacently. Cyanobacteria are proposed as good candidates for bioremediation of pollutants (Marchetto *et al.* 2021; Dudeja *et al.* 2025), and the result of this study indicates that some members may have evolved to cope with PFAS toxicity. The 14 *Microcystis* strains come from distinct areas and there is no apparent structure between sequence and geographical origin, which further supports a selection towards the observed genetic organizations or a potential ancestral genetic arrangement. All strains are from PFAS polluted waters at maximum concentration ranging from 5.3 to 1,450,000 ng/L (Saito *et al.* 2003; Choi *et al.* 2021; Evides 2024; Chang *et al.* 2025; Gao *et al.* 2025), except for strain NRERC-214 from the Caloosahatchee River in Florida, USA, for which PFAS pollution is seemingly not documented. This strain is also the only one without associated stress/photosynthesis genes adjacent to the FAcD gene, suggesting a lack of adaptation and supporting the selection for degradation and alleviation of photosynthesis inhibition.

The suspect FAcD is potentially ancestral to *Microcystis*, since it occurred in all 14 chromosomes of this genus in RefSeq complete genomes except for that of strain MC19, which is isolated from a lake in South Korea (Jeong *et al.* 2018) with no reported PFAS contamination. Furthermore, draft genomes were checked as well, using AllTheBacteria (Hunt *et al.* 2024) database version 0.1 (results not shown) and homologs of NOS0089 were found in all 8 available draft genomes of *Microcystis*. This supports a possible ancestral state of a FAcD homolog of NOS0089, although more genomes should be investigated and the hypothetical PFAS degradation potential of *Microcystis* should be tested.

Besides the 14 *Microcystis* genomes, homologs of NOS0089.1 are also found in 23 other complete bacterial genomes, of which cyanobacterium *Nostoc* and soil-associated *Skermanella* each is represented by 4 chromosomal regions (not shown). To a lesser degree than in *Microcystis*, the NOS0089.1 FAcD homologous gene is flanked by IS elements or stress-related functions. In the cyanobacterium *Stanieria cyanosphaera* PCC 7437 a gene encoding a PSII biogenesis protein Psp29 is adjacent to the FAcD homolog. The homolog is also found in the chromosomes of the phototrophic *Roseiflexus* (Hanada *et al.* 2002) and *Candidatus* Chlorohelix (Tsuji *et al.* 2024).

The other highly mobilized FacD BAE94252.1 and Q01398.1 (Fig. 2) matched homologs in the exact same RefSeq chromosomes (Supplementary Fig. S3). In four *Ralstonia* and two *Mesorhizobium* chromosomes, FAcD homologs are found adjacent to a gene encoding an uncharacterized putative thioesterase. The region surrounding the FAcD is more conserved in the included *Ralstonia* genomes, with several mobile genetic elements occurring in the vicinity, including multiple IS elements and a group II intron. Other nearby genes encode transcriptional regulators, propionate catabolism functions, and a gene potentially involved in cell cycle control (Supplementary Fig. S3).

## Conclusion and perspectives

This study provides a genomic survey of genes encoding potential PFAS-degrading enzymes in all complete bacterial genomes in the RefSeq database. Certain enzyme classes - particularly some defluorinating FAcDs and HADs - exhibit signs of mobilization and genetic synteny suggesting positive selection. Notably, the conserved presence of FAcD homologs in cyanobacteria, especially *Microcystis*, alongside genes involved in stress response, photosynthesis, and toxin-antitoxin systems, points to a possible evolutionary adaptation to PFAS-induced physiological stress rather than metabolic utilization. Our findings support a model in which microbial defluorination may evolve as a detoxification mechanism, particularly in environments where PFAS interfere with essential cellular processes such as membrane integrity and photosynthesis.

Future research should prioritize functional validation of these candidate enzymes, particularly those found in phototrophic and stress-adapted bacteria. Cloning and heterologous expression of FAcD and HAD variants, coupled with biochemical assays, will be critical to confirm their defluorination activity. Additionally, further studies are needed on the ecological contexts and selective pressures that favor the mobilization and fixation of genes encoding PFAS-degrading enzymes and associated genetic factors to alleviate stress and toxicity.

## Acknowledgements

This work was supported by Independent Research Fund Denmark [grant number 3105-00303B to T.K.N.].

## References

Altschul SF, Gish W, Miller W et al. Basic local alignment search tool. J Mol Biol 1990;215:403– 10.

Bhattacharya A, Fathima J, Varghese S et al. Advances in bioremediation strategies for PFAS-contaminated water and soil. Soil Environ Health 2025;3:100126.

Blin K. ncbi-genome-download. 2023, DOI: 10.5281/zenodo.8192486.

Buchfink B, Xie C, Huson DH. Fast and sensitive protein alignment using DIAMOND. Nat Methods 2015;12:59–60.

Bygd MD, Aukema KG, Richman JE et al. Unexpected Mechanism of Biodegradation and Defluorination of 2,2-Difluoro-1,3-Benzodioxole by Pseudomonas putida F1. mBio 2021;12, DOI: 10.1128/mBio.03001-21.

Chang W, Xu S-D, Liu T et al. Risk prioritization and experimental validation of per- and polyfluoroalkyl substances (PFAS) in Chaohu Lake: Based on nontarget and target analyses. J Hazard Mater 2025;492:138179.

Chetverikov S, Hkudaygulov G, Sharipov D et al. Biodegradation Potential of C7-C10 Perfluorocarboxylic Acids and Data from the Genome of a New Strain of Pseudomonas mosselii 5(3). Toxics 2023;11, DOI: 10.3390/toxics11121001.

Choi G-H, Lee D-Y, Bruce-Vanderpuije P et al. Environmental and dietary exposure of perfluorooctanoic acid and perfluorooctanesulfonic acid in the Nakdong River, Korea. Environ Geochem Health 2021;43:347–60.

Díaz-Orejas R, Espinosa M, Yeo CC. The Importance of the Expendable: Toxin–Antitoxin Genes in Plasmids and Chromosomes. Front Microbiol 2017;8, DOI: 10.3389/fmicb.2017.01479.

Dudeja C, Masroor S, Mishra V et al. Cyanobacteria-based bioremediation of environmental contaminants: advances and computational insights. Discov Agric 2025;3:42.

Edgar RC. Search and clustering orders of magnitude faster than BLAST. Bioinformatics 2010;26:2460–1.

Evides. PFAS en drinkwater. https://www.evides.nl/uw-drinkwater/mijn-waterkwaliteit/pfas-en-drinkwater (June 12, 2025,date last accessed)

Fenton SE, Ducatman A, Boobis A et al. Per- and Polyfluoroalkyl Substance Toxicity and Human Health Review: Current State of Knowledge and Strategies for Informing Future Research. Environ Toxicol Chem 2021;40:606–30.

Fitzgerald NJM, Simcik MF, Novak PJ. Perfluoroalkyl Substances Increase the Membrane Permeability and Quorum Sensing Response in Aliivibrio fischeri. Environ Sci Technol Lett 2018;5:26–31.

Fitzgerald NJM, Wargenau A, Sorenson C et al. Partitioning and Accumulation of Perfluoroalkyl Substances in Model Lipid Bilayers and Bacteria. Environ Sci Technol 2018;52:10433– 40.

Gao Y, Yang C, Feng G et al. Downward migration of per- and polyfluoroalkyl substances (PFAS) in lake sediments: Reconsideration of temporal trend analysis. J Hazard Mater 2025;492:138290.

Gilchrist CLM, Chooi Y-H. clinker & clustermap.js: automatic generation of gene cluster comparison figures. Bioinformatics 2021;37:2473–5.

Hanada S, Takaichi S, Matsuura K et al. Roseiflexus castenholzii gen. nov., sp. nov., a thermophilic, filamentous, photosynthetic bacterium that lacks chlorosomes. Int J Syst Evol Microbiol 2002;52:187–93.

Harris JD, Coon CM, Doherty ME et al. Engineering and characterization of dehalogenase enzymes from Delftia acidovorans in bioremediation of perfluorinated compounds. Synth Syst Biotechnol 2022;7:671–6.

Hu J, Wang D, Zhang N et al. Effects of perfluorooctanoic acid on Microcystis aeruginosa: Stress and self-adaptation mechanisms. J Hazard Mater 2023;445:130396.

Hu M, Scott C. Toward the development of a molecular toolkit for the microbial remediation of per-and polyfluoroalkyl substances. Nikel PI (ed.). Appl Environ Microbiol 2024;90:e00157–24.

Hualing M. Cyanobacterial NDH-1 Complexes. Front Microbiol 2022;13, DOI: 10.3389/fmicb.2022.933160.

Huang S, Jaffé PR. Defluorination of Perfluorooctanoic Acid (PFOA) and Perfluorooctane Sulfonate (PFOS) by Acidimicrobium sp. Strain A6. Environ Sci Technol 2019;53:11410– 9.

Hunt M, Lima L, Anderson D et al. AllTheBacteria - all bacterial genomes assembled, available and searchable. 2024:2024.03.08.584059.

Jaffé PR, Huang S, Park J et al. Defluorination of PFAS by Acidimicrobium sp. strain A6 and potential applications for remediation. Methods in Enzymology. Vol 696. Elsevier, 2024, 287–320.

Jeong H, Chun S-J, Srivastava A et al. Genome Sequences of Two Cyanobacterial Strains, Toxic Green Microcystis aeruginosa KW (KCTC 18162P) and Nontoxic Brown Microcystis sp. Strain MC19, under Xenic Culture Conditions. Genome Announc 2018;6:10.1128/genomea.00378-18.

Khusnutdinova AN, Batyrova KA, Brown G et al. Structural insights into hydrolytic defluorination of difluoroacetate by microbial fluoroacetate dehalogenases. FEBS J 2023;290:4966–83.

Klinkert B, Ossenbühl F, Sikorski M et al. PratA, a Periplasmic Tetratricopeptide Repeat Protein Involved in Biogenesis of Photosystem II in Synechocystis sp. PCC 6803 *. J Biol Chem 2004;279:44639–44.

Kojima H, Fukui M. Sulfuritalea hydrogenivorans gen. nov., sp. nov., a facultative autotroph isolated from a freshwater lake. Int J Syst Evol Microbiol 2011;61:1651–5.

Kolde R. pheatmap: Pretty Heatmaps. 2025.

Kong R, Xu X, Hu Z. A TPR-family membrane protein gene is required for light-activated heterotrophic growth of the cyanobacterium Synechocystis sp. PCC 6803. FEMS Microbiol Lett 2003;219:75–9.

Leong LEX, Khan S, Davis CK et al. Fluoroacetate in plants - a review of its distribution, toxicity to livestock and microbial detoxification. J Anim Sci Biotechnol 2017;8:55.

Liao J, Huang L, Liu Y et al. Metabolomic analysis reveals contrasting effects of PFOS and PFAS on cyanobacterial bloom and metabolic pathways in eutrophic water. Environ Pollut 2025;375:126340.

Lindell AE, Grießhammer A, Michaelis L et al. Extensive PFAS accumulation by human gut bacteria. 2024:2024.09.17.613493.

Liu G, Zhang S, Yang K et al. Toxicity of perfluorooctane sulfonate and perfluorooctanoic acid to Escherichia coli: Membrane disruption, oxidative stress, and DNA damage induced cell inactivation and/or death. Environ Pollut 2016;214:806–15.

Liu J, Mejia Avendaño S. Microbial degradation of polyfluoroalkyl chemicals in the environment: A review. Environ Int 2013;61:98–114.

Liu X, Li Y, Zheng X et al. Anti-oxidant mechanisms of Chlorella pyrenoidosa under acute GenX exposure. Sci Total Environ 2021;797:149005.

Liu Z, Cao X, Wu M et al. Mechanisms of PFBA toxicity in Chlorella vulgaris: Photosynthesis, oxidative stress, and antioxidant impairment. Environ Res 2025;273:121228.

Marchetto F, Roverso M, Righetti D et al. Bioremediation of Per-and Poly-Fluoroalkyl Substances (PFAS) by Synechocystis sp. PCC 6803: A Chassis for a Synthetic Biology Approach. Life 2021;11:1300.

Murphy CD, Schaffrath C, O’Hagan D. Fluorinated natural products: the biosynthesis of fluoroacetate and 4-fluorothreonine in Streptomyces cattleya. Chemosphere 2003;52:455–61.

Naumann A, Alesio J, Poonia M et al. PFAS fluidize synthetic and bacterial lipid monolayers based on hydrophobicity and lipid charge. J Environ Chem Eng 2022;10:107351.

Néron B, Littner E, Haudiquet M et al. IntegronFinder 2.0: Identification and Analysis of Integrons across Bacteria, with a Focus on Antibiotic Resistance in Klebsiella. Microorganisms 2022;10:700.

Nielsen TK, Browne PD, Hansen LH. Antibiotic resistance genes are differentially mobilized according to resistance mechanism. GigaScience 2022;11:giac072.

Nielsen TK, Rasmussen M, Demanèche S et al. Evolution of Sphingomonad Gene Clusters Related to Pesticide Catabolism Revealed by Genome Sequence and Mobilomics of Sphingobium herbicidovorans MH. Genome Biol Evol 2017;9:2477–90.

Nielsen TK, Xu Z, Gözdereliler E et al. Novel Insight into the Genetic Context of the cadAB Genes from a 4-chloro-2-methylphenoxyacetic Acid-Degrading Sphingomonas. PLOS ONE 2013;8:e83346.

Oksanen J. vegan: community ecology package. R Package Version 2015;2:3.

Parsons JR, Sáez M, Dolfing J et al. Biodegradation of Perfluorinated Compounds. In: Whitacre DM (ed.), Reviews of Environmental Contamination and Toxicology Vol 196. New York, NY: Springer US, 2008, 53–71.

Pearce SL, Oakeshott JG, Pandey G. Insights into Ongoing Evolution of the Hexachlorocyclohexane Catabolic Pathway from Comparative Genomics of Ten Sphingomonadaceae Strains. G3 GenesGenomesGenetics 2015;5:1081–94.

Presentato A, Lampis S, Vantini A et al. On the ability of perfluorohexane sulfonate (PFHxS) bioaccumulation by two Pseudomonas sp. strains isolated from PFAS-contaminated environmental matrices. Microorganisms 2020;8, DOI: 10.3390/microorganisms8010092.

Qiao W, Xie Z, Zhang Y et al. Perfluoroalkyl substances (PFASs) influence the structure and function of soil bacterial community: Greenhouse experiment. Sci Total Environ 2018;642:1118–26.

Rosendahl S, Tamman H, Brauer A et al. Chromosomal toxin-antitoxin systems in Pseudomonas putida are rather selfish than beneficial. Sci Rep 2020;10:9230.

Ross K, Varani AM, Snesrud E et al. TnCentral: a Prokaryotic Transposable Element Database and Web Portal for Transposon Analysis. mBio 2021;12:10.1128/mbio.02060-21.

Saito N, Sasaki K, Nakatome K et al. Perfluorooctane Sulfonate Concentrations in Surface Water in Japan. Arch Environ Contam Toxicol 2003;45:149–58.

Salinero KK, Keller K, Feil WS et al. Metabolic analysis of the soil microbe Dechloromonas aromatica str. RCB: indications of a surprisingly complex life-style and cryptic anaerobic pathways for aromatic degradation. BMC Genomics 2009;10:351.

Schwengers O, Jelonek L, Dieckmann MA et al. Bakta: Rapid and standardized annotation of bacterial genomes via alignment-free sequence identification. Microb Genomics 2021;7, DOI: 10.1099/MGEN.0.000685,.

Shaw DMJ, Munoz G, Bottos EM et al. Degradation and defluorination of 6:2 fluorotelomer sulfonamidoalkyl betaine and 6:2 fluorotelomer sulfonate by Gordonia sp. strain NB4-1Y under sulfur-limiting conditions. Sci Total Environ 2019;647:690–8.

Siguier P, Perochon J, Lestrade L et al. ISfinder: the reference centre for bacterial insertion sequences. Nucleic Acids Res 2006;34:D32–6.

Smorada CM, Sima MW, Jaffé PR. Bacterial degradation of perfluoroalkyl acids. Curr Opin Biotechnol 2024;88:103170.

Sperfeld M, Diekert G, Studenik S. Anaerobic aromatic compound degradation in Sulfuritalea hydrogenivorans sk43H. FEMS Microbiol Ecol 2019;95:fiy199.

Stockbridge RB, Wackett LP. The link between ancient microbial fluoride resistance mechanisms and bioengineering organofluorine degradation or synthesis. Nat Commun 2024;15:4593.

Tsuji JM, Shaw NA, Nagashima S et al. Anoxygenic phototroph of the Chloroflexota uses a type I reaction centre. Nature 2024;627:915–22.

Wackett LP. Why Is the Biodegradation of Polyfluorinated Compounds So Rare? McMahon K (ed.). mSphere 2021;6:e00721–21.

Wackett LP. Confronting PFAS persistence: enzymes catalyzing C–F bond cleavage. Trends Biochem Sci 2025;50:71–83.

Wang P, Duan W, Takabayashi A et al. Chloroplastic NAD(P)H Dehydrogenase in Tobacco Leaves Functions in Alleviation of Oxidative Damage Caused by Temperature Stress. Plant Physiol 2006;141:465–74.

Wang Z, Dewitt JC, Higgins CP et al. A Never-Ending Story of Per- and Polyfluoroalkyl Substances (PFASs)? Environ Sci Technol 2017;51:2508–18.

Weathers TS, Higgins CP, Sharp JO. Enhanced Biofilm Production by a Toluene-Degrading Rhodococcus Observed after Exposure to Perfluoroalkyl Acids. Environ Sci Technol 2015;49:5458–66.

Wickham H. Reshaping Data with the reshape Package. J Stat Softw 2007;21:1–20.

Wickham H. dplyr: A grammar of data manipulation. R Package Version 04 2015;3:p156.

Wickham H. ggplot2 -Elegant Graphics for Data Analysis (2nd Edition). J Stat Softw 2016;77, DOI: 10.18637/jss.v077.b02.

Wickham H. Tidy Messy Data://CRAN. R-Project. Org/Package= Tidyr. https://tidyr.tidyverse.org/ (June 12, 2025, xdate last accessed)

Wijayahena MK, Moreira IS, Castro PML et al. PFAS biodegradation by Labrys portucalensis F11: Evidence of chain shortening and identification of metabolites of PFOS, 6:2 FTS, and 5:3 FTCA. Sci Total Environ 2025;959:178348.

Wintenberg ME, Vasilyeva OB, Schaffter SW. Comparative Transcriptomic Analysis of Perfluoroalkyl Substances-Induced Responses of Exponential and Stationary Phase Escherichia coli. 2025:2025.02.18.638913.

Yang M, Ye J, Qin H et al. Influence of perfluorooctanoic acid on proteomic expression and cell membrane fatty acid of Escherichia coli. Environ Pollut 2017;220:532–9.

Yi LB, Chai LY, Xie Y et al. Isolation, identification, and degradation performance of a PFOA-degrading strain. Genet Mol Res 2016;15, DOI: 10.4238/gmr.15028043.

