## Supplemental data for "Mobilization of genes encoding potential PFAS-degradation enzymes and positive selection in cyanobacteria"

Contents

- Supplementary Table 1. Enzymes used in this study for finding homologous sequencens.
- Supplementary Figure S1. Amino acid similarities between target enzymes.
- Supplementary Figure S2. Compression rate of clustering hit regions to Clustered Hit Loci (CHLs).
- Supplementary Fig. S3. Genetic context of FAcD BAE94252.1 homologs in *Ralstonia* and *Mesorhizobium* chromosomes.

Supplementary Table 1. Enzymes used in this study for finding homologous sequencens.

| **Accession number (and other accession number found in litterature)** | **Product and strain of origin** | **MOBscale** | **Occur-rences** | **References, substrates, and notes** | **Enzyme class** |
|---|---|---|---|---|---|
| BAE94252.1 (Q1JU72) | MenH FAc-DEX fluoroacetate dehalogenase [Burkholderia sp. FA1] | 0.54 | 6 | Substrate: fluoroacetate (Kurihara *et al*. 2003; Jitsumori *et al*. 2009; Hu and Scott 2024) | Fluoroacetate dehalogenase |
| Q01398.1 | DehH1 H-1 Haloacetate dehalogenase [Delftia acidovorans] | 0.54 | 6 | Substrate: Fluoroacetate (Sota *et al*. 2002; Hu and Scott 2024)<br>Has been described as a HAD but is actually a FAcD based on sequence analysis (has catalytic triad) and is placed on FAcD branch in tree. | Fluoroacetate dehalogenase |
| WP_012052601.1 | TodA toluene dioxygenase alpha-subunit [Pseudomonas putida F1] | 0.51 | 52 | Substrate: 2,2-Difluoro-1,3-Benzodioxole. The full operon is *todABC1C2* (Bygd *et al*. 2021) | Dioxygenase |
| AB970510.1 | BtfD 2-hydroxy-6-oxo-7,7,7-trifluorohepta-2,4-dienoate hydrolase [Rhodococcus sp. 065240] | 0.46 | 170 | Substrate: Aromatic fluorinated compounds (Yano *et al*. 2015) | Dioxygenase |
| PZP66635.1 | haloacid dehalogenase [Delftia | 0.44 | 14 | Substrate: PFOA, although not significantly | Haloacid dehalogenase |

| | | | | | |
|---|---|---|---|---|---|
| | acidovorans MAG] | | | different from negative control (Harris *et al*. 2022; Hu and Scott 2024). | |
| NOS0089 | Alpha/beta hydrolase fold protein [Nostoc sp.] | 0.39 | 35 | Substrate: difluoroacetate (Khusnutdinova *et al*. 2023) | Haloacid dehalogenase |
| CAA12245.1 | OAH 6-oxocyclohex-1-ene-1-carbonyl-CoA hydrolase [Thauera aromatica] | 0.37 | 19 | Substrate: 2-fluorobenzoate (Tiedt *et al*. 2017) | Hydrolase |
| CAA12246.1 | DCH cyclohexa-1,5-diene-1-carbonyl-CoA hydratase [Thauera aromatica] | 0.36 | 17 | Substrate: 2-fluorobenzoate (Tiedt *et al*. 2017) | Hydratase |
| WP_275630221.1 | AlkB-2 alkane 1-monooxygenase [Pseudomonas sp. 273] | 0.35 | 44 | Substrate: fluorinated alkanes. Genes discovered by transcriptomics but not verified with cloning (Xie *et al*. 2023) | Monooxygenase |
| X79076.1_cbdB | CbdB 2-halobenzoate 1,2-dioxygenase [P. cepacia 2CBS] | 0.35 | 127 | Substrate: 2-halobenzoates (Haak, Fetzner and Lingens 1995) | Dioxygenase |
| WP_238404007.1 (RS16300) | Alcohol dehydrogenase catalytic domain-containing protein [Gordonia sp. NB41Y] | 0.35 | 71 | Substrate: potentially 6:2 FTAB, based on transcriptomics (Bottos *et al*. 2020) | Dehydrogenase |
| WP_020793433.1 (RS22855) | LLM class flavin- | 0.33 | 788 | Substrate: potentially 6:2 | Oxidoreductase |

| | | | | | |
|---|---|---|---|---|---|
| | dependent oxidoreductase [Gordonia sp. NB41Y] | | | FTAB and 6:2 FTSA, based on transcriptomics (Bottos *et al*. 2020) | |
| POL0530 (ABE42492.1, Bpro0530) | haloacid dehalogenase, type II [Polaromonas sp. JS666] | 0.33 | 2 | Substrate: fluoroacetate (Chan *et al*. 2022a; Khusnutdinova *et al*. 2023; Hu and Scott 2024) | Haloacid dehalogenase |
| BAL41322.1 | Maleylacetate reductase [Ralstonia pickettii DTP0602] | 0.32 | 112 | Substrate: 2,4,6-trichlorophenol and 4-fluorobenzoate (Hatta, Fujii and Takizawa 2012) | Reductase |
| WP_053776268.1 (RS02605) | NAD(P)-dependent alcohol dehydrogenase [Gordonia sp. NB41Y] | 0.32 | 2140 | Substrate: potentially 6:2 FTAB and 6:2 FTSA, based on transcriptomics (Bottos *et al*. 2020) | Dehydrogenase |
| AAD55885.1 | MacA hypothetical protein [Cupriavidus necator 335T] | 0.31 | 25 | Substrate: 4-fluorobenzoate (Seibert *et al*. 2004) | Dehydrogease |
| DeHa4 | Fluoroacetate dehalogenase [Delftia acidovorans D4B] | 0.30 | 647 | Substrate: di- and monofluoroacetate (Farajollahi *et al*. 2024) | Fluoroacetate dehalogenase |
| WP_269074981.1 | SsuD Alkanesulfonate monooxygenase/LLM class flavin-dependent oxidoreductase [Dietzia aurantiaca] | 0.29 | 727 | Substrate: potentially 6:2 FTSA (Méndez *et al*. 2022), although evidence might be incomplete (Hu and Scott 2024) | Oxidoreductase |

| | | | | | |
|---|---|---|---|---|---|
| MCD2262844.1 | SsuD LLM class flavin-dependent oxidoreductase [Dietzia aurantiaca] | 0.29 | 735 | Substrate: potentially 6:2 FTSA (Méndez *et al*. 2022) , although evidence might be incomplete (Hu and Scott 2024) | Oxidoreductase |
| WP_053776025.1 (RS00415) | LLM class flavin-dependent oxidoreductase [Gordonia sp. NB41Y] | 0.29 | 456 | Substrate: potentially 6:2 FTAB and 6:2 FTSA, based on transcriptomics (Bottos *et al*. 2020) | Oxidoreductase |
| WP_053777181.1 (RS10415) | LLM class flavin-dependent oxidoreductase [Gordonia sp. NB41Y] | 0.28 | 41 | Substrate: potentially 6:2 FTAB, 6:2 FTSA, and octanesulfonate based on transcriptomics (Bottos *et al*. 2020) | Oxidoreductase |
| RJO0230 (Rha0230) | Haloacid dehalogenase [Rhodococcus jostii strain RHA1] | 0.28 | 22 | Substrate: fluoroacetate, difluoroacetate, 6:2 FTOH, and 6:2 FTSA (Chan *et al*. 2022b; Khusnutdinova *et al*. 2023; Hu and Scott 2024) | Haloacid dehalogenase |
| X79076.1_cbdA | 2-halobenzoate 1,2-dioxygenase [Pseudomonas cepacia 2CBS] | 0.27 | 577 | Subtrate: fluorobenzoate (Haak, Fetzner and Lingens 1995) | Dioxygenase |
| WP_053777000.1 (RS08865) | Zinc-binding dehydrogenase [Gordonia sp. NB41Y] | 0.26 | 84 | Substrate: potentially 6:2 FTAB and octanesulfonate based on transcriptomics | Dehydrogenase |

| | | | | | |
|---|---|---|---|---|---|
| | | | | (Bottos *et al.* 2020) | |
| ABG92213.1 | cytochrome P450 CYP254 [Rhodococcus jostii RHA1] | 0.26 | 38 | Substrate: 6:2 FTOH based on transcriptomics but needs further evidence (Yang *et al.* 2022) | Cytochrome P450 |
| WP_053777622.1 (RS14165) | LLM class flavin-dependent oxidoreductase [Gordonia sp. NB41Y] | 0.25 | 60 | Substrate: potentially 6:2 FTAB, 6:2 FTSA, and octanesulfonate based on transcriptomics (Bottos *et al.* 2020) | Oxidoreductase |
| ABG94339.1 | alkane 1-monooxygenase [Rhodococcus jostii RHA1] | 0.25 | 337 | Substrate: 6:2 FTOH based on transcriptomics but needs further evidence (Yang *et al.* 2022) | Monooxygenase |
| AAX84119.1 | putative benzoyl-CoA reductase, partial [Thauera aromatica] | 0.24 | 25 | Substrate: 4-fluorobenzoate and 4-fluorotoluene (Tiedt *et al.* 2016) | Reductase |
| CAD15064.1 (RSc1362) | putative dehalogenase-like hydrolase; protein [Ralstonia pseudosolanacearum GMI1000] | 0.24 | 225 | No defluorination activity on fluoroacetate (Chan *et al.* 2022b) | Haloacid dehalogenase |
| AAX84174.1 | putative benzoyl-CoA reductase, partial [Thauera aromatica] | 0.22 | 25 | Substrate: 4-fluorobenzoate and 4-fluorotoluene (Tiedt *et al.* 2016) | Reductase |
| AAX84166.1 | putative benzoyl-CoA reductase, partial | 0.22 | 25 | Substrate: 4-fluorobenzoate and 4-fluorotoluene | Reductase |

| | | | | | |
|---|---|---|---|---|---|
| | [Thauera aromatica] | | | (Tiedt *et al.* 2016) | |
| BAI96793.1 (K01563) | linB/dhaA haloalkane dehalogenase [Sphingobium japonicum UT26S] | 0.21 | 89 | Substrate: dehalogenation of linuron but no detected defluorination (Dong *et al.* 2024) | Haloalkane dehalogenase |
| AAG04199.1 (PA0810) | probable haloacid dehalogenase [Pseudomonas aeruginosa PAO1] | 0.17 | 168 | No defluorination activity on fluoroacetate (Chan *et al.* 2022b) | Haloacid dehalogenase |
| RPA1163 (3R3U_1) | Fluoroacetate dehalogenase [Rhodopseudomonas palustris] | 0.15 | 145 | Substrate: fluoroacetate, difluoroacetate, chloroacetate, 2-fluoropropionic acid, 2,3,3,3-tetrafluoropropionic acids (Khusnutdinova *et al.* 2023; Hu and Scott 2024). Substrate promiscuity: several compounds including phenyl and benzyls (Hu and Scott 2024). | Haloacid dehalogenase |
| WP_053777639.1 (RS14160) | LLM class flavin-dependent oxidoreductase [Gordonia sp. NB41Y] | 0.12 | 11 | Substrate: potentially 6:2 FTAB and 6:2 FTSA, based on transcriptomics (Bottos *et al.* 2020) | Oxidoreductase |
| WP_275629779.1 | alkB-3 alkane 1-monooxygenase | 0.07 | 180 | Substrate: 1,10-difluorodecane based on transcriptomics (Xie *et al.* 2023) | Monooxygenase |

| | | | | | |
|---|---|---|---|---|---|
| | [Pseudomonas sp. 273] | | | | |
| WP_053778136.1 (RS18475) | NADP-dependent oxidoreductase [Gordonia sp. NB41Y] | 0.04 | 8 | Substrate: potentially 6:2 FTAB and 6:2 FTSA, based on transcriptomics (Bottos *et al*. 2020) | Oxidoreductase |
| DAR3835 | Alpha/beta hydrolase fold protein [Dechloromonas aromatica] | 0.00 | 1 | Substrate: Fluoroacetate, difluoroacetate, 2,2-difluoropropionic acid, 5,5,5-trifluoropentanoic acid (Khusnutdinova *et al*. 2023; Hu and Scott 2024) | Haloacid dehalogenase |
| AXG51384.1 | LmbB2 peroxygenase-like enzyme [Streptomyces lincolnensis LC-G] | 0.00 | 1 | Substrate: 3-fluoro-L-tyrosine (Wang *et al*. 2019) | Peroxygenase |
| WP_053777670.1 (RS14725) | LLM class flavin-dependent oxidoreductase [*Gordonia* sp. NB41Y] | 0.00 | 1 | Substrate: potentially 6:2 FTAB and octanesulfonate based on transcriptomics (Bottos *et al*. 2020) | Oxidoreductase |
| WP_011137954.1 (DeHa2) | haloacid dehalogenase type II [*Delftia acidovorans*] | NA | 0 | Substrate: Defluorination of mono- and difluoroacetate (Harris *et al*. 2022; Farajollahi *et al*. 2024) | Haloacid dehalogenase |
| QAX87819.1 | incomplete putative haloacid dehalogenase (RdhA) from [*Acidimicrobia* | NA | 0 | Substrate: PFOA and PFOS. Knockout mutants lose the ability to | Haloacid dehalogenase |

| | | | | |
|---|---|---|---|---|
| | *ceae bacterium* A6] | | | defluorinate (Jaffé *et al.* 2024) |

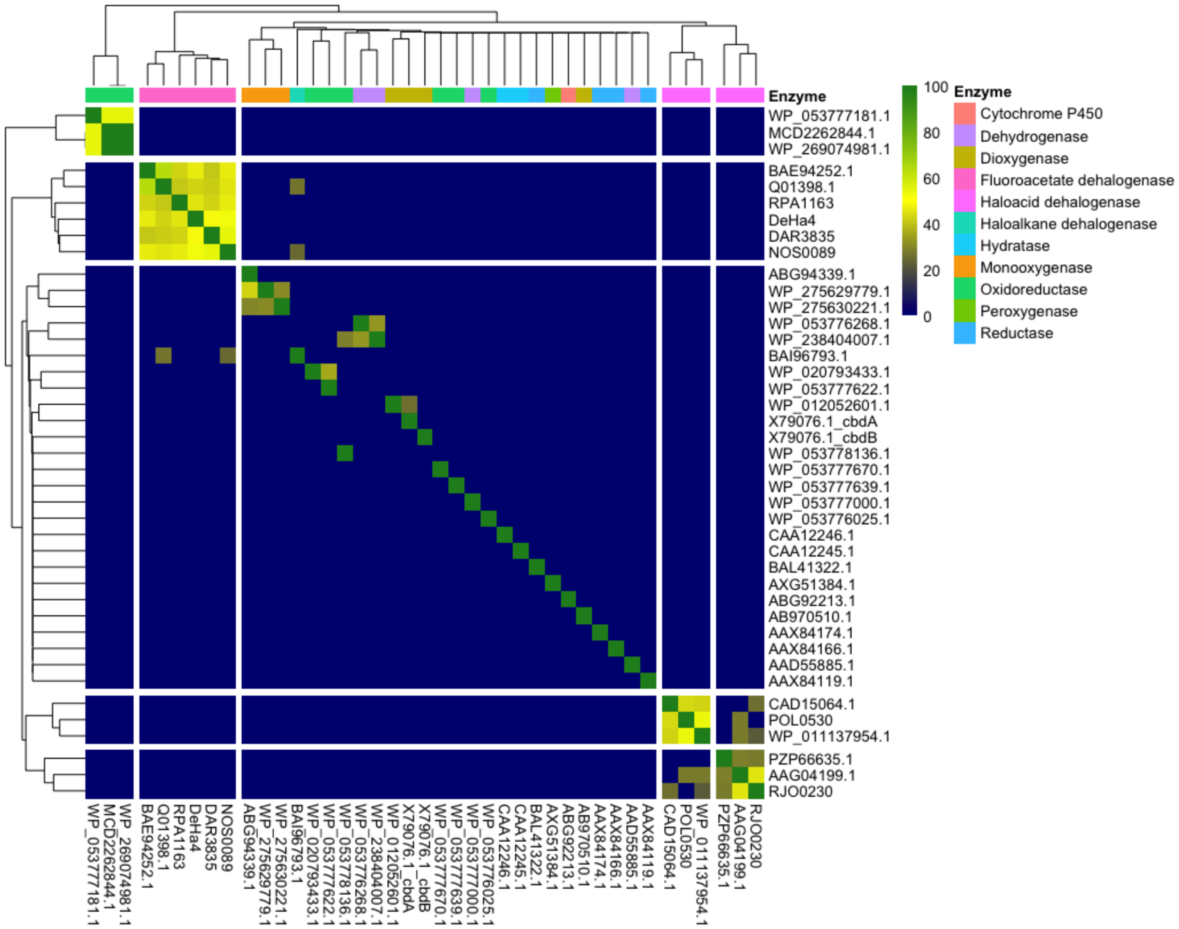

Supplementary Fig. S1. Heatmap of BLASTP similarities of selected defluorinating enzymes. Columns and rows are clustered with hierarchical clustering, as implemented in the pheatmap package. Only BLASTP hits with a minimum E-value of 10E-10 and query coverage of >80% are shown.

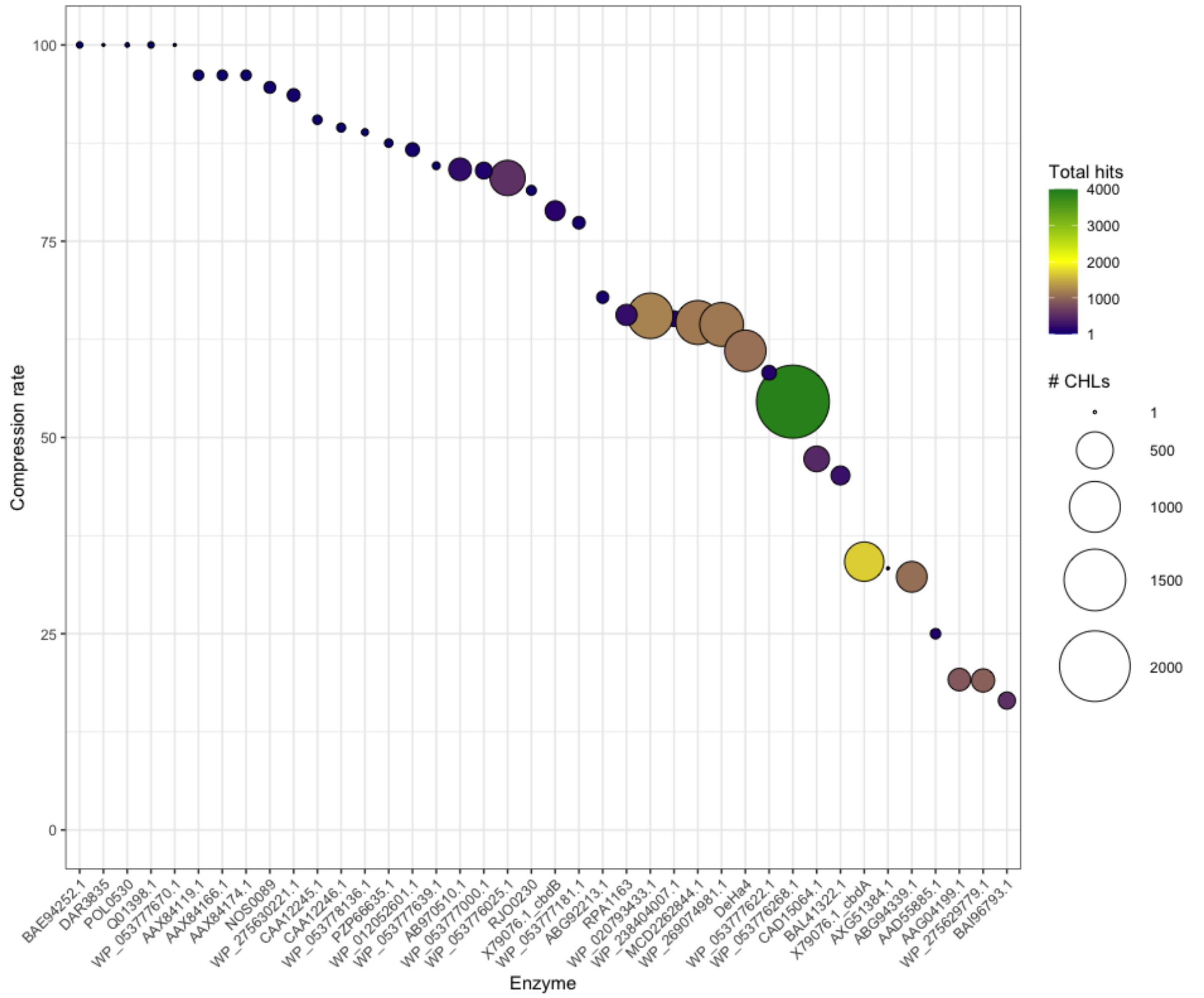

*Supplementary Fig. S2. Compression rate of clustering hit regions to Clustered Hit Loci (CHLs). Hit regions include the target gene plus 12,170 bp flanking regions on each side. Clustering was performed with USEARCH. The compression rate is expressed as number of clusters (CHLs) divided by total hits for a given gene.*

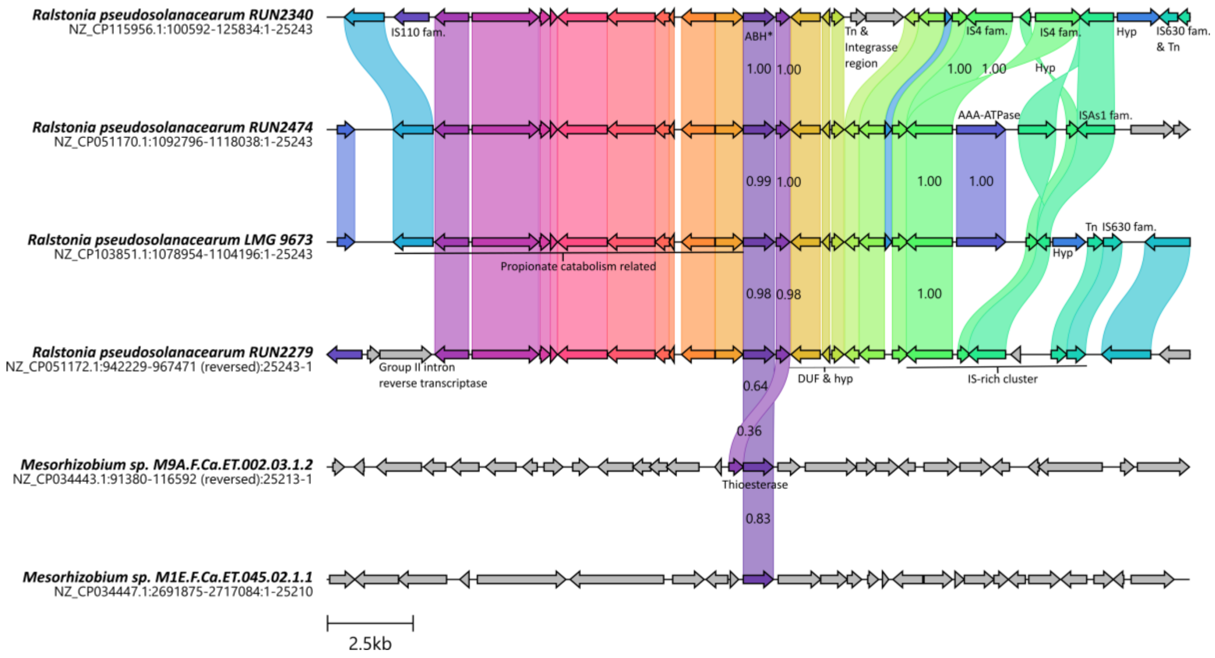

Supplementary Fig. S3. Genetic context of FAcD BAE94252.1 homologs in Ralstonia and Mesorhizobium chromosomes. ABH is abbreviation for alpha/beta hydrolase. Numbers in links indicate similarity between amino acid sequences of similar genes. Annotations were manually curated to highlight genes of interest, including IS elements and propionate catabolism. The figure was produced with Clinker and manually curated in InkScape.

**Supplementary references**

Bottos EM, AL-shabib EY, Shaw DMJ *et al.* Transcriptomic response of Gordonia sp. strain NB4-1Y when provided with 6:2 fluorotelomer sulfonamidoalkyl betaine or 6:2 fluorotelomer sulfonate as sole sulfur source. *Biodegradation* 2020;**31**:407–22.

Bygd MD, Aukema KG, Richman JE *et al.* Unexpected Mechanism of Biodegradation and Defluorination of 2,2-Difluoro-1,3-Benzodioxole by Pseudomonas putida F1. *mBio* 2021;**12**, DOI: 10.1128/mBio.03001-21.

Chan PWY, Chakrabarti N, Ing C *et al.* Defluorination Capability of l-2-Haloacid Dehalogenases in the HAD-Like Hydrolase Superfamily Correlates with Active Site Compactness. *ChemBioChem* 2022a;**23**:e202100414.

Chan PWY, Chakrabarti N, Ing C *et al.* Defluorination Capability of l-2-Haloacid Dehalogenases in the HAD-Like Hydrolase Superfamily Correlates with Active Site Compactness. *ChemBioChem* 2022b;**23**:e202100414.

Dong S, Yan PF, Mezzari MP *et al.* Using Network Analysis and Predictive Functional Analysis to Explore the Fluorotelomer Biotransformation Potential of Soil Microbial Communities. *Environ Sci Technol* 2024;**58**:7480–92.

Farajollahi S, Lombardo NV, Crenshaw MD *et al.* Defluorination of Organofluorine Compounds Using Dehalogenase Enzymes from Delftia acidovorans (D4B). *ACS Omega* 2024;**9**:28546–55.

Haak B, Fetzner S, Lingens F. Cloning, nucleotide sequence, and expression of the plasmid-encoded genes for the two-component 2-halobenzoate 1,2-dioxygenase from Pseudomonas cepacia 2CBS. *J Bacteriol* 1995;**177**:667–75.

Harris JD, Coon CM, Doherty ME *et al.* Engineering and characterization of dehalogenase enzymes from Delftia acidovorans in bioremediation of perfluorinated compounds. *Synth Syst Biotechnol* 2022;**7**:671–6.

Hatta T, Fujii E, Takizawa N. Analysis of Two Gene Clusters Involved in 2,4,6-Trichlorophenol Degradation by Ralstonia pickettii DTP0602. *Biosci Biotechnol Biochem* 2012;**76**:892–9.

Hu M, Scott C. Toward the development of a molecular toolkit for the microbial remediation of per-and polyfluoroalkyl substances. Nikel PI (ed.). *Appl Environ Microbiol* 2024;**90**:e00157-24.

Jaffé PR, Huang S, Park J *et al.* Defluorination of PFAS by Acidimicrobium sp. strain A6 and potential applications for remediation. *Methods in Enzymology*. Vol 696. Academic Press Inc., 2024, 287–320.

Jitsumori K, Omi R, Kurihara T *et al.* X-Ray Crystallographic and Mutational Studies of Fluoroacetate Dehalogenase from Burkholderia sp. Strain FA1. *J Bacteriol* 2009;**191**:2630–7.

Khusnutdinova AN, Batyrova KA, Brown G *et al.* Structural insights into hydrolytic defluorination of difluoroacetate by microbial fluoroacetate dehalogenases. *FEBS J* 2023;**290**:4966–83.

Kurihara T, Yamauchi T, Ichiyama S *et al.* Purification, characterization, and gene cloning of a novel fluoroacetate dehalogenase from *Burkholderia* sp. FA1. *J Mol Catal B Enzym* 2003;**23**:347–55.

Méndez V, Holland S, Bhardwaj S *et al.* Aerobic biotransformation of 6:2 fluorotelomer sulfonate by *Dietzia aurantiaca* J3 under sulfur-limiting conditions. *Sci Total Environ* 2022;**829**:154587.

Seibert V, Thiel M, Hinner I-S *et al.* Characterization of a gene cluster encoding the maleylacetate reductase from Ralstonia eutropha 335T, an enzyme recruited for growth with 4-fluorobenzoate. *Microbiology* 2004;**150**:463–72.

Sota M, Endo M, Nitta K *et al.* Characterization of a Class II Defective Transposon Carrying Two Haloacetate Dehalogenase Genes from Delftia acidovorans Plasmid pUO1. *Appl Environ Microbiol* 2002;**68**:2307–15.

Tiedt O, Mergelsberg M, Boll K *et al.* ATP-Dependent C–F Bond Cleavage Allows the Complete Degradation of 4-Fluoroaromatics without Oxygen. *mBio* 2016;**7**, DOI: 10.1128/mBio.00990-16.

Tiedt O, Mergelsberg M, Eisenreich W *et al.* Promiscuous defluorinating enoyl-CoA hydratases/hydrolases allow for complete anaerobic degradation of 2-fluorobenzoate. *Front Microbiol* 2017;**8**, DOI: 10.3389/fmicb.2017.02579.

Wang Y, Davis I, Shin I *et al.* Biocatalytic Carbon-Hydrogen and Carbon-Fluorine Bond Cleavage through Hydroxylation Promoted by a Histidyl-Ligated Heme Enzyme. *ACS Catal* 2019;**9**:4764–76.

Xie Y, Ramirez D, Chen G *et al.* Genome-Wide Expression Analysis Unravels Fluoroalkane Metabolism in Pseudomonas sp. Strain 273. *Environ Sci Technol* 2023;**57**:15925–35.

Yang SH, Shi Y, Strynar M *et al.* Desulfonation and defluorination of 6:2 fluorotelomer sulfonic acid (6:2 FTSA) by Rhodococcus jostii RHA1: Carbon and sulfur sources, enzymes, and pathways. *J Hazard Mater* 2022;**423**:127052.

Yano K, Wachi M, Tsuchida S *et al.* Degradation of benzotrifluoride via the dioxygenase pathway in Rhodococcus sp. 065240. *Biosci Biotechnol Biochem* 2015;**79**:496–504.
